# Variation Benchmark Datasets: Update, Criteria, Quality and Applications

**DOI:** 10.1101/634766

**Authors:** Anasua Sarkar, Yang Yang, Mauno Vihinen

**Author notes:** Correspondence to Mauno Vihinen, Department of Experimental Medical Science, BMC B13, Lund University, SE-22 184 Lund, Sweden.

## Abstract

Development of new computational methods and testing their performance has to be done on experimental data. Only in comparison to existing knowledge can method performance be assessed. For that purpose, benchmark datasets with known and verified outcome are needed. High-quality benchmark datasets are valuable and may be difficult, laborious and time consuming to generate. VariBench and VariSNP are the two existing databases for sharing variation benchmark datasets. They have been used for training and benchmarking predictors for various types of variations and their effects. There are 419 new datasets from 109 papers containing altogether 329003373 variants; however there is plenty of redundancy between the datasets. VariBench is freely available at http://structure.bmc.lu.se/VariBench/. The contents of the datasets vary depending on information in the original source. The available datasets have been categorized into 20 groups and subgroups. There are datasets for insertions and deletions, substitutions in coding and non-coding region, structure mapped, synonymous and benign variants. Effect-specific datasets include DNA regulatory elements, RNA splicing, and protein property predictions for aggregation, binding free energy, disorder and stability. Then there are several datasets for molecule-specific and disease-specific applications, as well as one dataset for variation phenotype effects. Variants are often described at three molecular levels (DNA, RNA and protein) and sometimes also at the protein structural level including relevant cross references and variant descriptions. The updated VariBench facilitates development and testing of new methods and comparison of obtained performance to previously published methods. We compared the performance of the pathogenicity/tolerance predictor PON-P2 to several benchmark studies, and showed that such comparisons are feasible and useful, however, there may be limitations due to lack of provided details and shared data.

**AUTHOR SUMMARY:** A prediction method performance can only be assessed in comparison to existing knowledge. For that purpose benchmark datasets with known and verified outcome are needed. High-quality benchmark datasets are valuable and may be difficult, laborious and time consuming to generate. We collected variation datasets from literature, website and databases. There are 419 separate new datasets, which however contain plenty of redundancy. VariBench is freely available at http://structure.bmc.lu.se/VariBench/. There are datasets for insertions and deletions, substitutions in coding and non-coding region, structure mapped, synonymous and benign variants. Effect-specific datasets include DNA regulatory elements, RNA splicing, and protein property predictions for aggregation, binding free energy, disorder and stability. Then there are several datasets for molecule-specific and disease-specific applications, as well as one dataset for variation phenotype effects. The updated VariBench facilitates development and testing of new methods and comparison of obtained performance to previously published methods. We compared the performance of the pathogenicity/tolerance predictor PON-P2 to several benchmark studies and showed that such comparisons are possible and useful when the details of studies and the datasets are shared.

## INTRODUCTION

Development and testing of computational methods are dependent on experimental data. Only in comparison to existing knowledge can method performance be assessed. For that purpose, benchmark datasets with known and verified outcome are needed. During the last few years, such datasets have been collected for a number of applications in the field of variation interpretation. VariBench [1] and VariSNP [2] are the two existing databases for variation benchmark datasets. VariBench contains all kinds of datasets while VariSNP is a dedicated resource for variation sets from dbSNP database for short variations [3].

Benchmark datasets are used both for method training and testing. We can divide testing approaches into three categories (Figure 1). The most reliable are systematic benchmark studies. Quite often the initial method performance assessment is done on somewhat limited test data or not reporting all necessary measures. The third group includes studies for initial method and hypothesis testing typically with a limited amount of data. An example for this kind of testing is Critical Assessment of Genome Interpretation (CAGI, https://genomeinterpretation.org/), which has organized several challenges for method developers. These contests with blind data, when the participants do not know the true answer, have been important e.g. for testing new ideas and methods, as well for tackling novel application areas.

**Figure 1.**
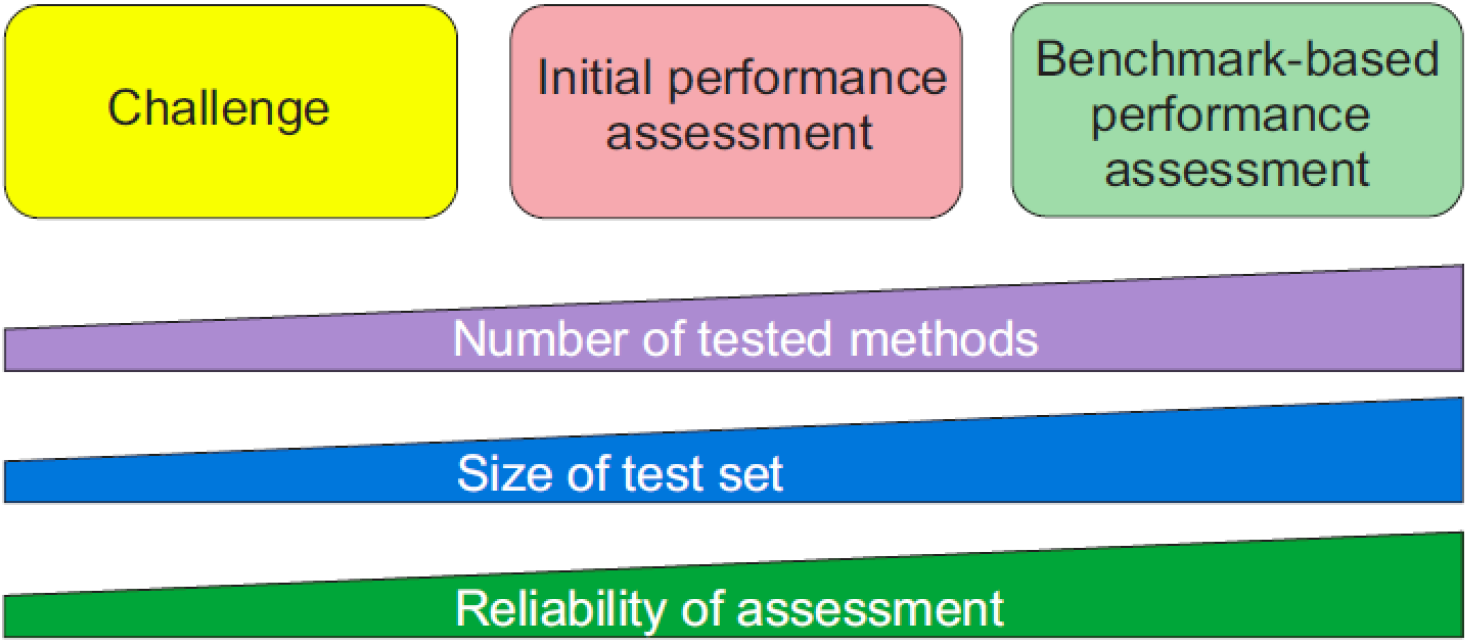
Types of method performance tests. The figure is adapted from [34].

High-quality benchmark datasets are valuable and may be difficult, laborious and time consuming to generate. Already from the point of view of reasonable use of resources it is important to share such datasets. Secondly, comparison of method performance is reliable only when using the same test dataset. According to the FAIR principles [4], research data should be made findable, accessible, interoperable and re-usable. VariBench and VariSNP provide variation data according to these principles.

It is still quite common that authors collect and use extensive datasets for their published papers, but do not share and make the datasets available. This prevents others from comparing additional tools to those used in the paper. Even when the data is made available, it may be in a format that makes re-use practically impossible. An example is the datasets used for testing the MutationTaster2 tolerance predictor [5]. They were published as figures and at barely legible resolution. Now, these datasets are available in VariBench.

## CRITERIA FOR BENCHMARKS

We defined criteria for a benchmark when the VariBench database was first published [1]. These criteria were more extensive than previously used and have been found very useful and still form the basis for inclusion of data and for their representation in VariBench. The criteria are as follows.

### Relevance

The dataset has to capture the characteristics of the investigated property. Not all available data may be relevant for the phenomenon or may be only indirectly related to it. The collected cases have to be for the specific effect or mechanism under study.

### Representativeness

The datasets should cover the event space as well as possible, thus preferably containing examples from all the regions relevant to the effect. The actual number of cases for achieving this coverage may vary widely depending on the effect. The dataset should be of sufficient size to allow statistical studies but may not need to include all known instances.

### Non-redundancy

This means excluding overlapping cases.

### Experimentally verified cases

Method performance comparisons have to be based on experimental data, not on predictions, otherwise the comparison will be about the congruence of methods, not about their true performance.

### Positive and negative cases

Comprehensive assessment has to be based both on positive (showing the investigated feature) and negative (not having effect) cases.

### Scalability

It should be possible to test systems of different sizes.

### Reusability

As datasets are expensive to generate they should be shared in such a way that they can be used for other investigations. This may mean similar applications or usage in new areas.

Most of the criteria are rather easy to fulfil, but some others are more difficult to take into account. We recently investigated the representativeness of 24 tolerance datasets from VariBench in the human protein universe by analysing the distribution and coverage of cases in chromosomes, protein structures, CATH domains and classes, Pfam families, Enzyme Commission (EC) categories and Gene Ontology annotations [6]. The outcome was that none of the datasets were well representative. When correlating the training data representativeness to the performance of predictors based on them, no clear correlation was found. However, it is apparent that representative training data would allow training of methods that have good performance for cases distributed throughout the event space.

Benchmark studies in relation to variation predictions have been made for variants affecting protein stability [7, 8], protein substitution tolerance/pathogenicity [9–14], protein localization [15], protein disorder [16], protein solubility [17], benign variants [18], transmembrane proteins [19], alternative splicing [20, 21] and phenotypes of amino acid substitutions [22]. Many of the datasets used in these studies are available for verification and reuse, but unfortunately e.g. the last one, which is unique, is not accessible.

To test relevance of the tolerance datasets, we investigated how many disease-causing variations could be found from neutral training data. A small number of such variants were found, 1.13 to 1.77 % [6]. These numbers are so small that they do not have a major impact on method performances. VariBench datasets are reusable and scalable, contain experimental cases, and are typically non-redundant. However, how redundancy should be defined may depend on the application. For example, when using domain features in variant predictors, variants even in related domain members would be redundant.

## DATASET QUALITY

The quality of benchmark datasets is of utmost significance. This is naturally dependent on the quality of the data sources. There are not many quality schemes in this field. For locus specific variation databases (LSDBs) there is a quality scheme that contains close to 50 criteria in four main areas including database quality, technical quality, accessibility and timeliness [23]. However, these guidelines are not yet widely followed and similar criteria are missing for other types of variation data resources.

Systematics within datasets and databases can significantly improve their quality and usability. For variation data there are a number of systematics solutions available. These include systematic gene names available for human from the HUGO Gene Nomenclature Committee (HGNC) [24], Human Genome Variation Society (HGVS) variation nomenclature [25], Locus Reference Genomic (LRG) and [26] RefSeq reference sequences [27], and Variation Ontology (VariO) variation type, effect and mechanism annotations [28].

Quality relates to numerous aspects in the datasets, the correctness of variation and gene/protein and disease information, relevance of references, etc. We recently selected cases from ProTherm [29] to build an unbiased dataset for the protein variant stability predictor PON-tstab [30]. We were aware that the database had some problems, however, were surprised with the extent of problematic cases. While making the selection, we noticed numerous issues, such as cases of two-stage denaturation pathways where values for all the steps and then the total value were provided; there were errors in sequences, variants, recorded measuring temperatures, ΔΔG values and their signs and units, and in indicated PDB structures; and so on. The uncorrected and wrong data have been used for development of tens of prediction methods. This is probably an extreme exception (ProTherm was taken away from the internet after our paper was published); however, this indicates that one has to be careful even when using popular data. VariBench has several quality controls, but lists also datasets that may contain problems e.g. numerous ProTherm sub-selections that have been published and sometimes used in several papers. They are included for comparative purposes.

## HOW TO TEST PREDICTOR PERFORMANCE

The use of a benchmark dataset is just one of the requirements for systematic method performance assessment. Proper measures are needed to find out the qualities of performance. Most of the currently available prediction methods are binary, distributing cases into two categories. There are guidelines for how to test and report method performance [31–33]. There is also a checklist what to report when using such methods in publications.

Results for binary methods are presented in a contingency (also called for confusion) table out of which different measures can be calculated. The most important ones are the following six, which according to the guidelines [32] have to be provided for comprehensive assessments. Specificity, sensitivity, positive and negative predictive values (PPV and NPV) use half of the data in the matrix, while accuracy and Matthews correlation coefficient (MCC) use data from all the four data cells. Additional useful measures include area under curve (AUC) when presenting Receiver Operating Characteristic (ROC) curves, and Overall Performance Measure (OPM). Good methods display a balanced performance, their values for measures differ only slightly.

In case there is an imbalance in the number of cases in the classes, it has to be mitigated [31]. Several approaches are available for that. Cases used for testing method performance should not have been used for training them, otherwise there is circularity that overinflates performance measures [14]. A scheme has been presented on how datasets should be split for training and testing as well as blind testing [34]. When there are more than two predicted classes additional measures are available [31, 32]. In addition to these measures, method assessment can contain other factors such as time required for predictions, as well as user friendliness and clarity of the service and results.

Datasets used for assessment have to be of sufficient size. There are a number of reasons for this requirement. Widely used machine learning methods are statistical by nature and require a relatively large number of cases for reliable testing. If we think the event space, in the case of proteins, there are 380 different amino acid substitution types, 150 of which are more likely due to happening because of a single nucleotide substitution within the coding region for a codon. These substitutions can appear in numerous different contexts, thus too small test datasets should be avoided. There are several performance assessments, especially for variants in a single protein or a small number of genes/proteins that do not have any statistical power. The smallest dataset we have seen contained just nine substitutions based on which a detailed analysis was performed to recommend the best performing tools!

Variation interpretation is often done in relation to human diseases. It is important to note that diseases are not binary states (benign/disease) instead there is a continuum and certain disease state can appear due to numerous different combinations of disease components, see the pathogenicity model [35]. This aspect has not been taken into account in benchmark datasets apart from training data for PON-PS [36] and clinical data for cystic fibrosis [37].

## VARIATION DATASETS

We have collected from literature, websites and databases datasets, which have been used for training and benchmarking various types of variations and their effects (Table 1). The new datasets come from 109 papers. There are 419 new separate datasets containing altogether 329003373 variants. One paper can contain more than one dataset. The number of unique variants is smaller as many of the datasets are different subsets of commonly used datasets such as ClinVar or ProTherm, or VariBench itself. The total number is dominated by VariSNP cases.

**Table 1.**
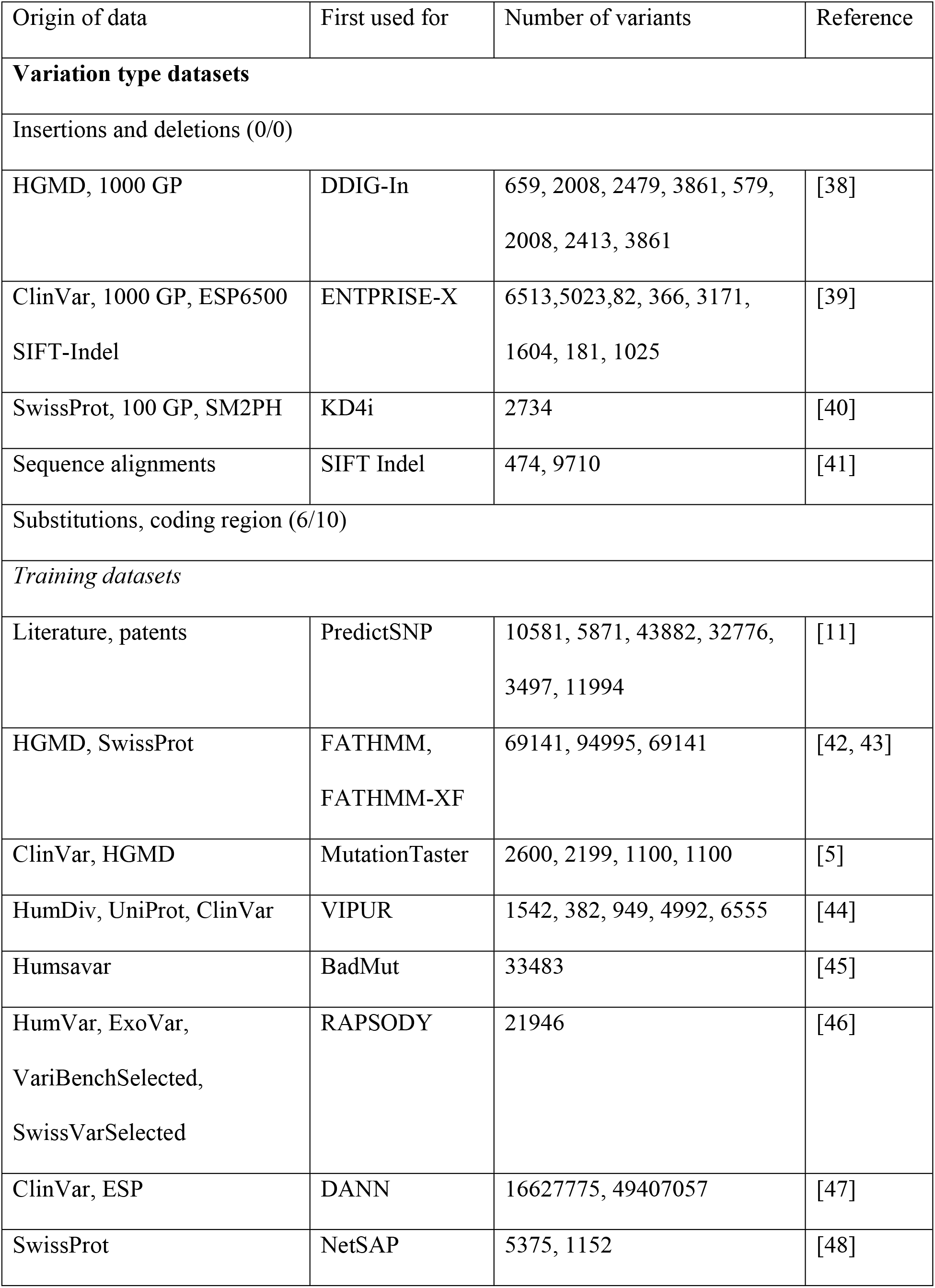
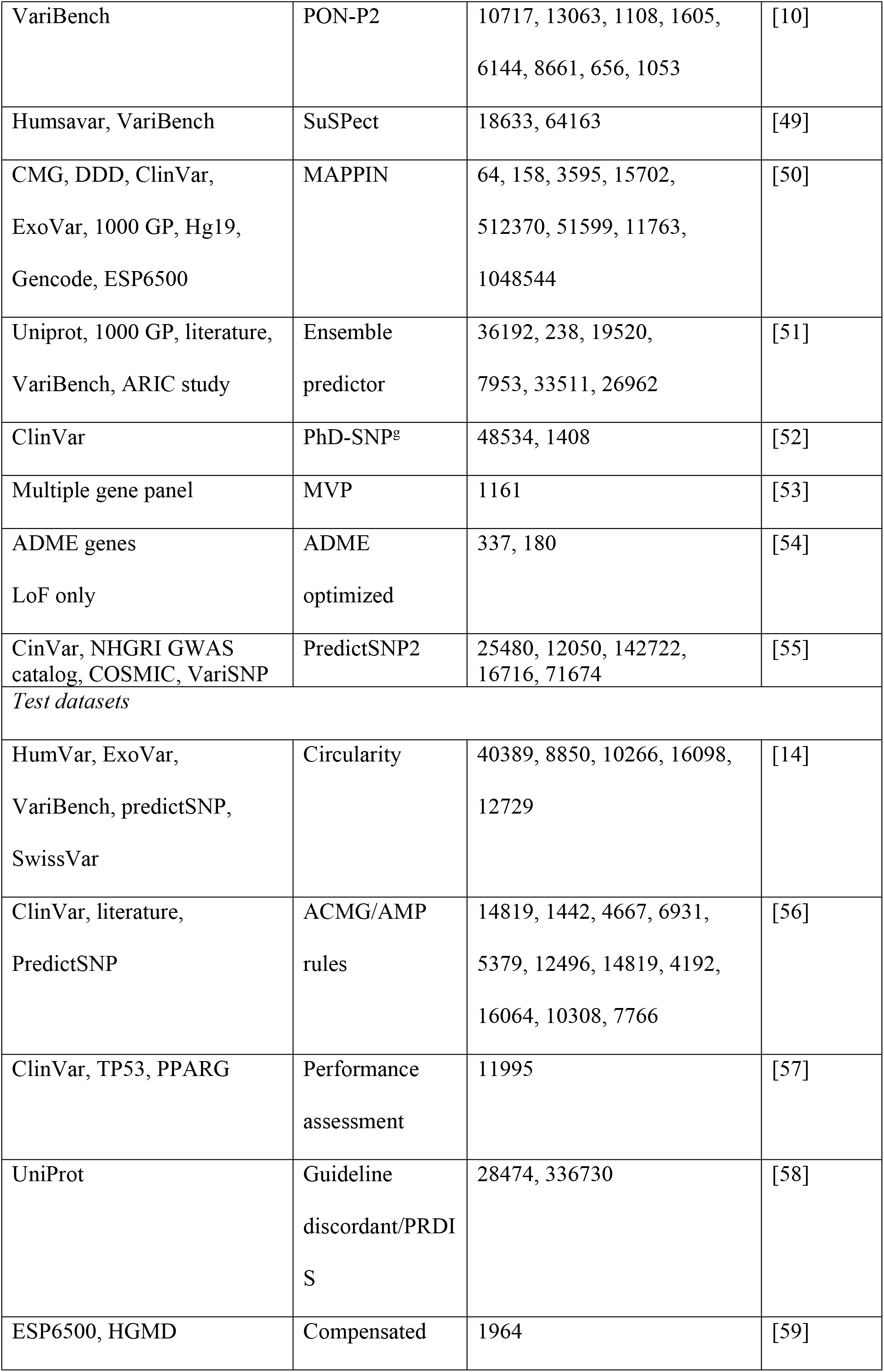
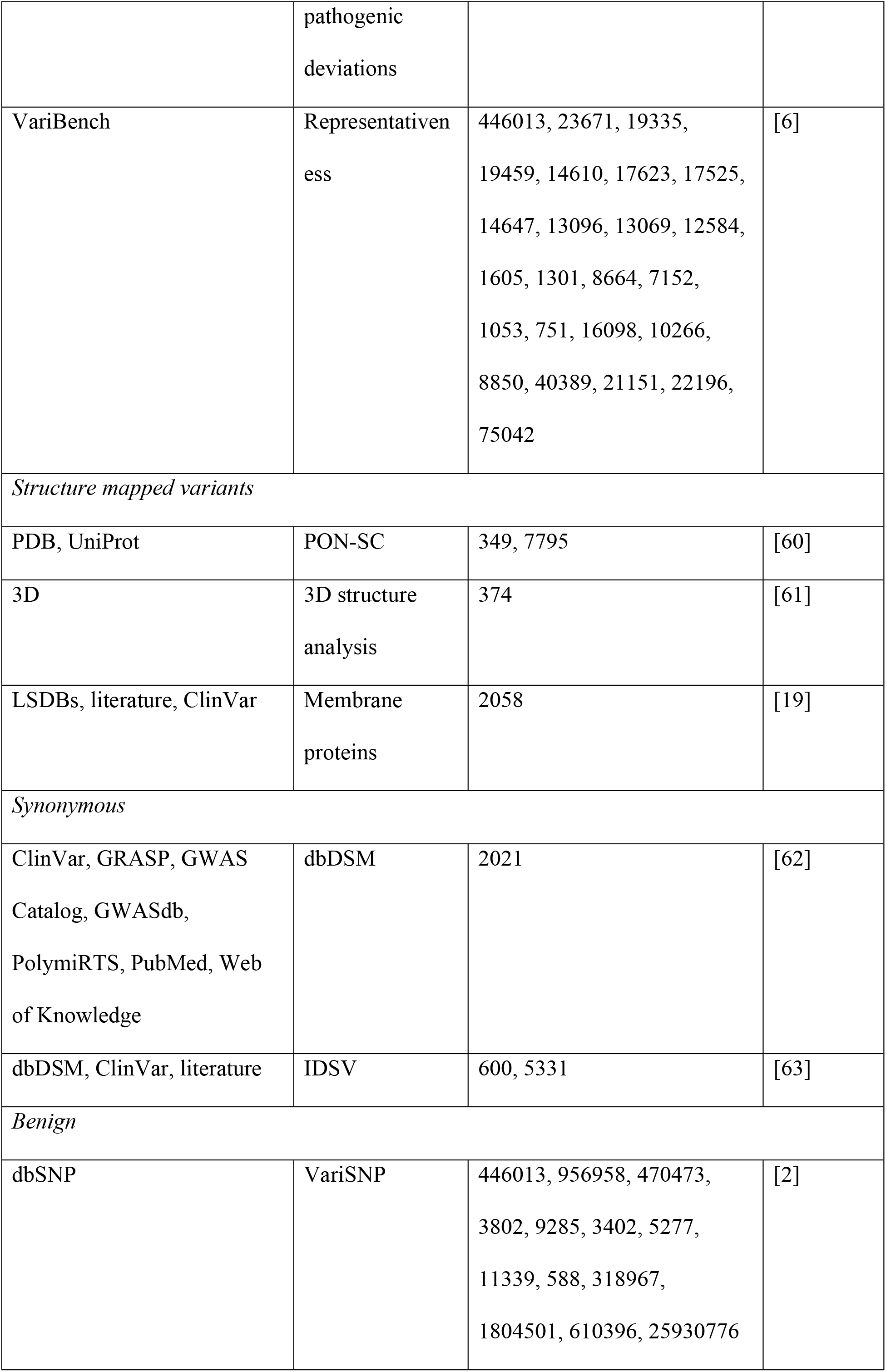
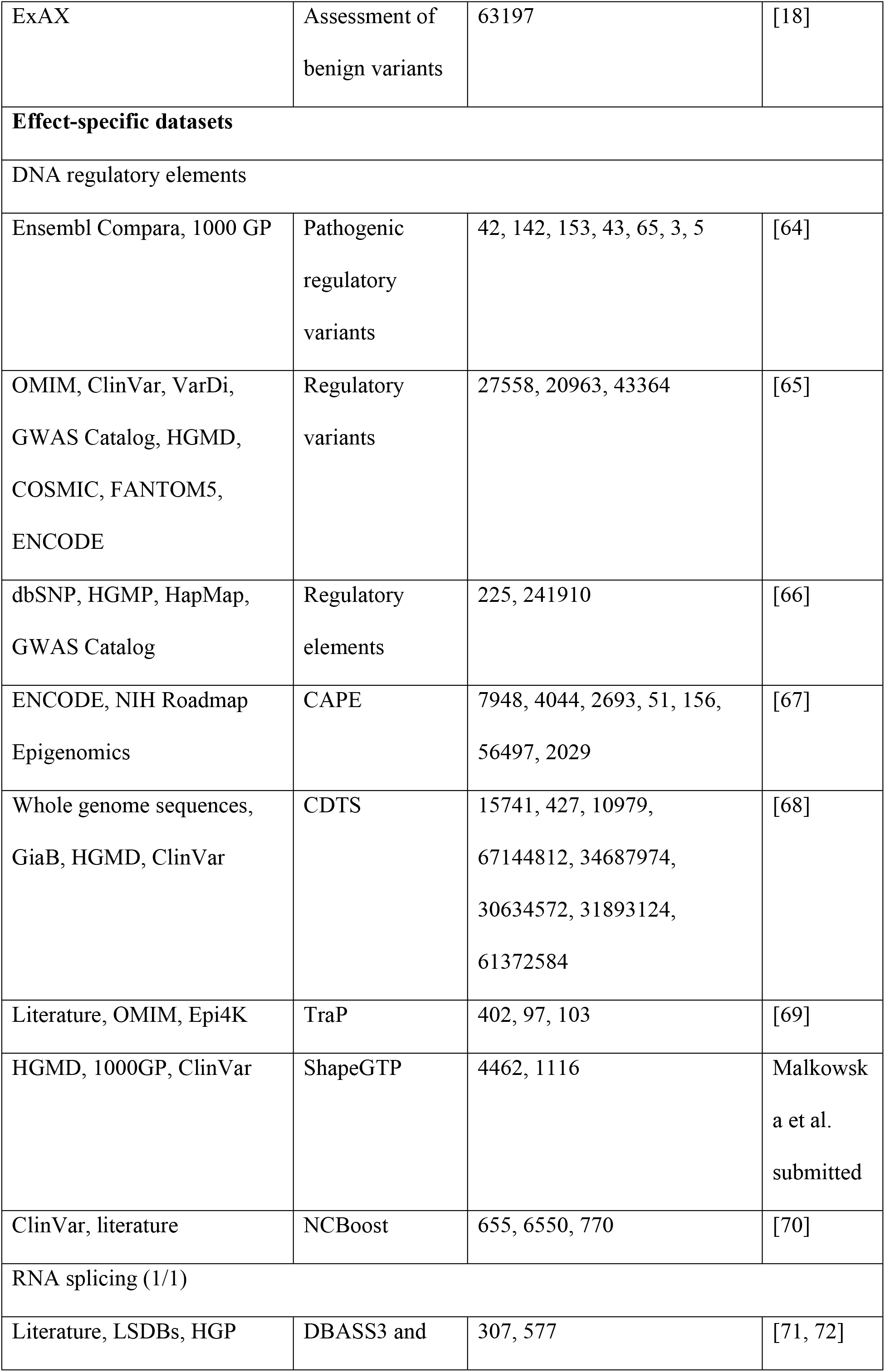
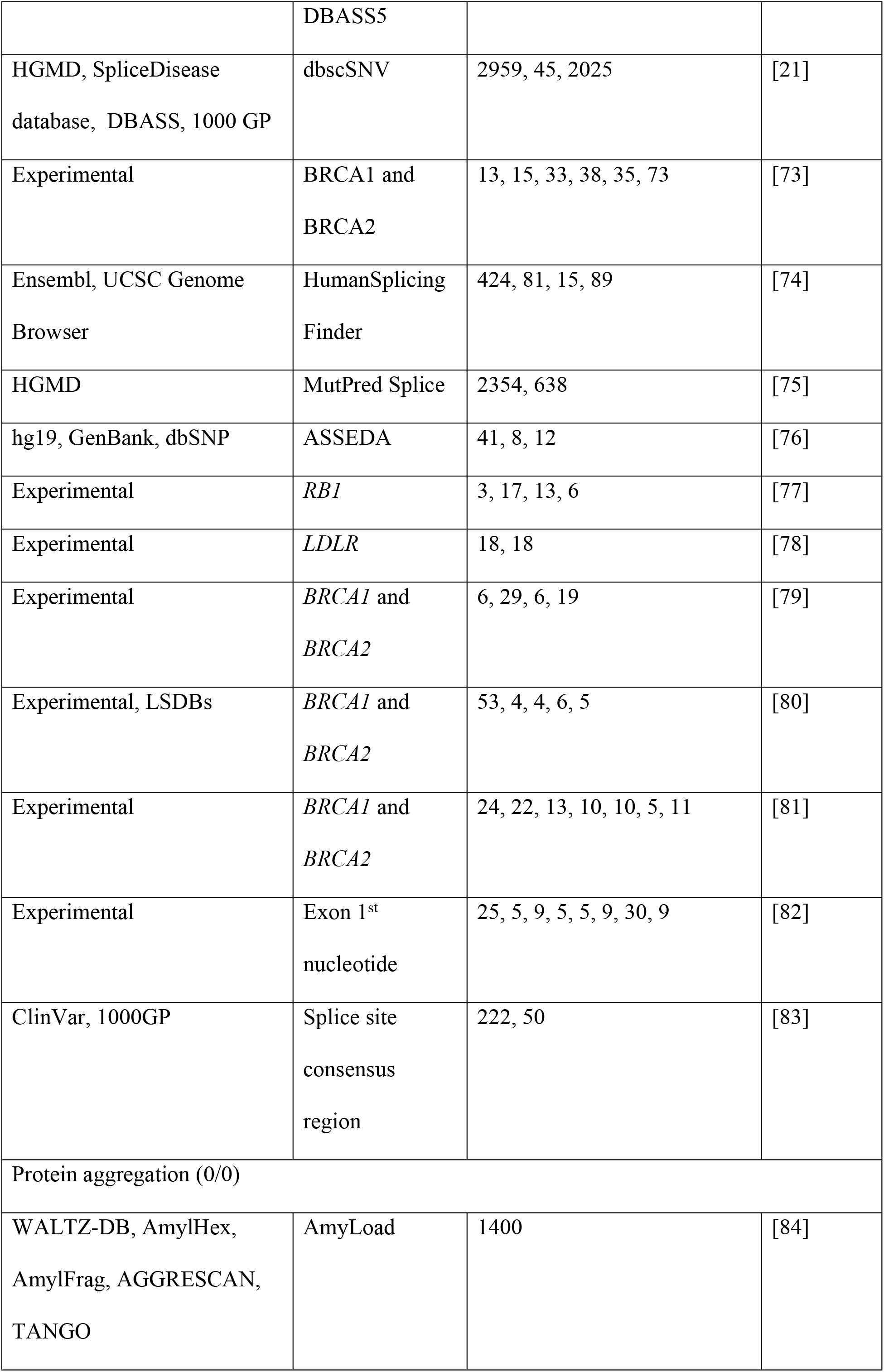
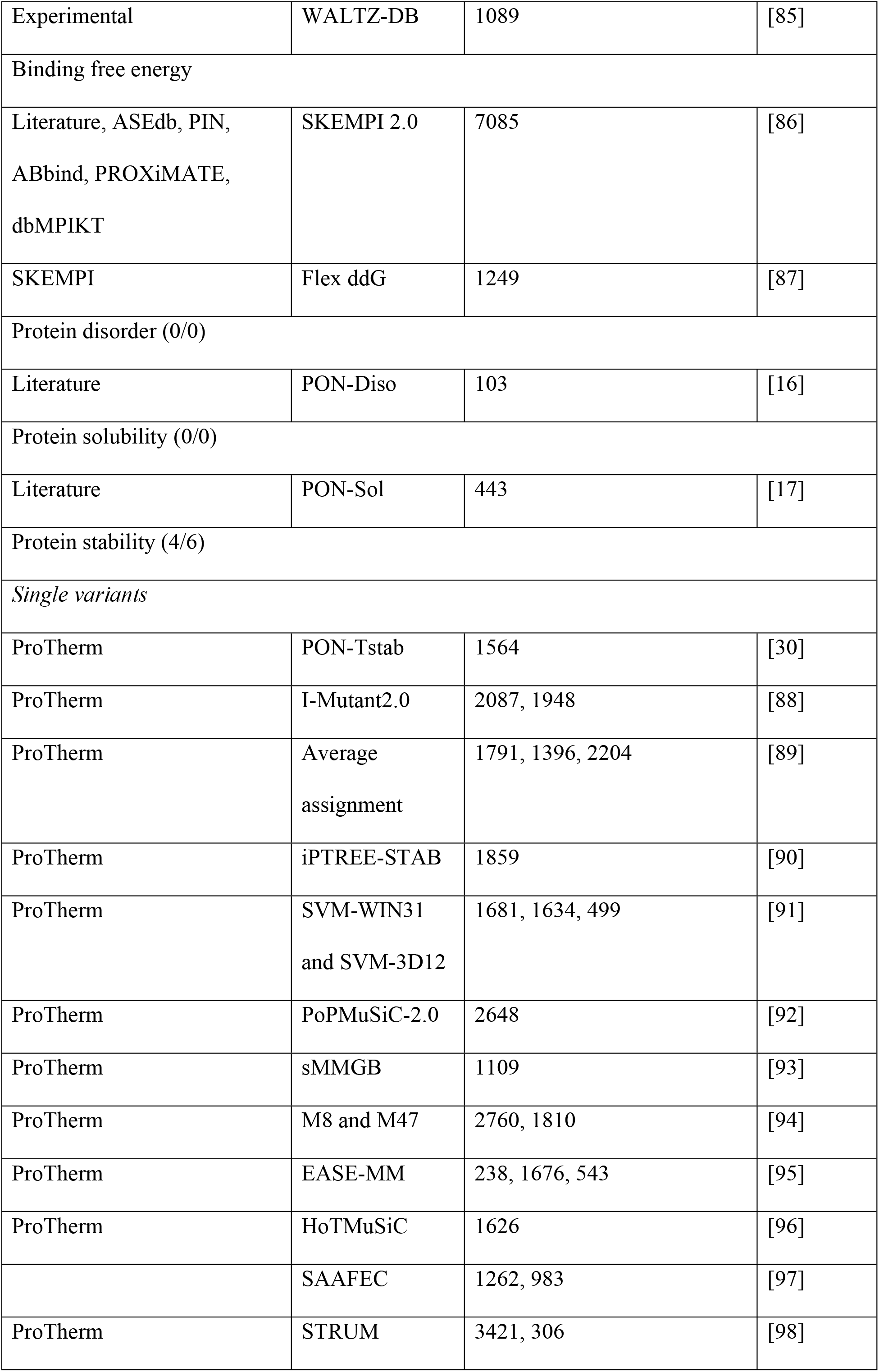
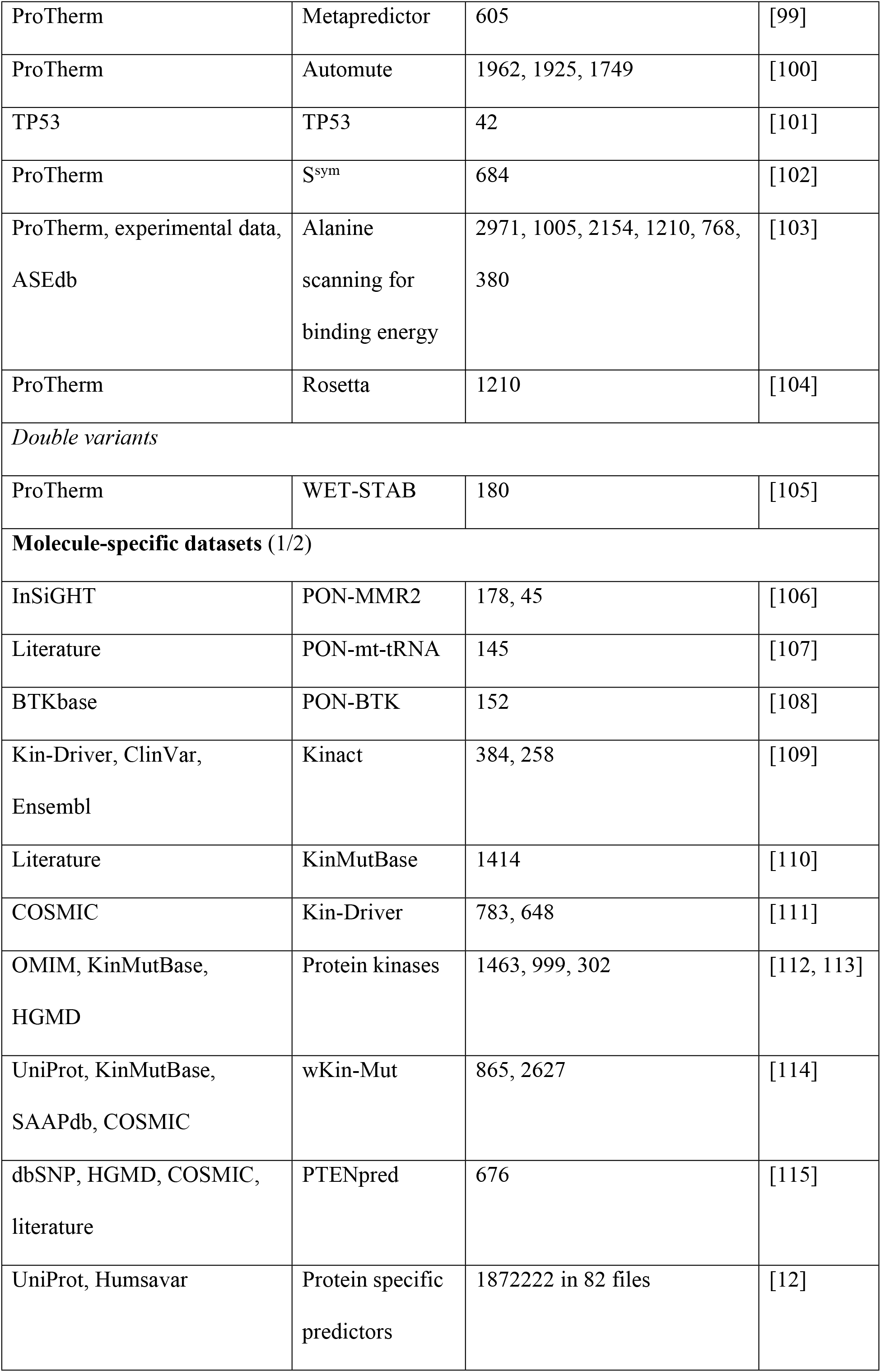
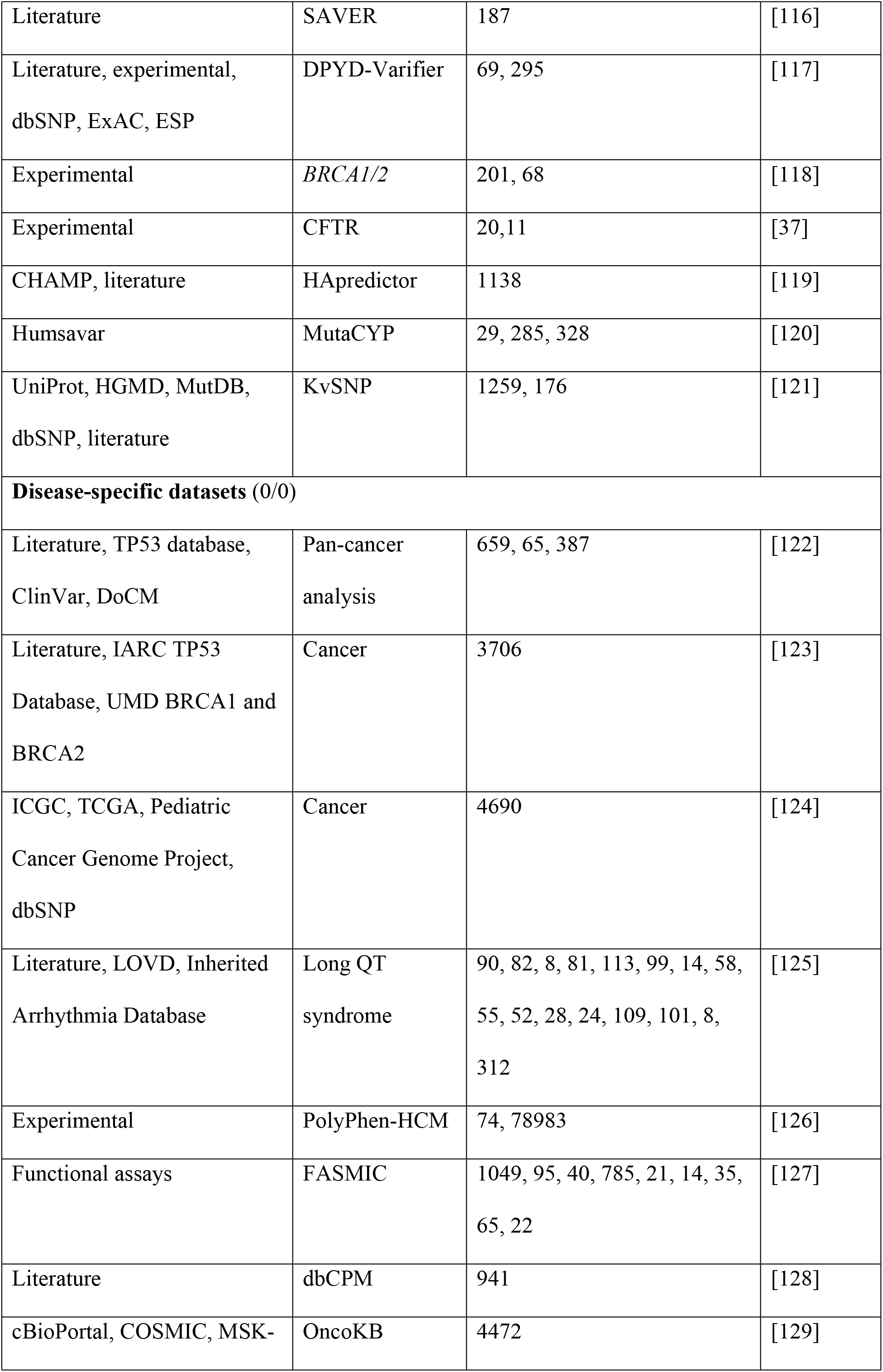
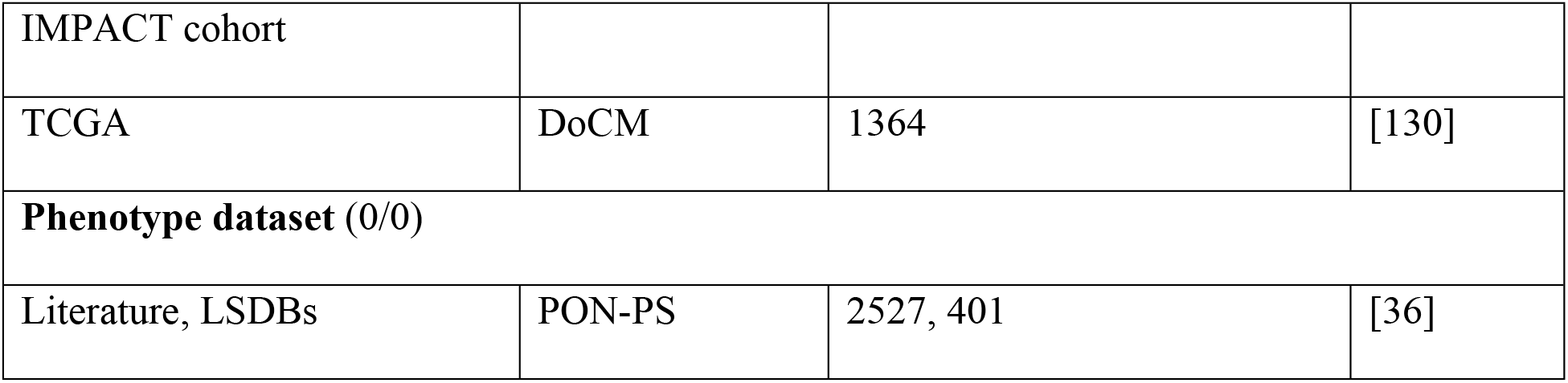
New benchmark datasets added to VariBench

VariBench datasets are freely available at http://structure.bmc.lu.se/VariBench/ and can be downloaded separately. The website contains basic information about the datasets, their origin and for what purpose they were initially used for. Datasets are categorized similar to Table 1 for easy access. The contents of the datasets vary depending on information in the original source. We have enriched many of them e.g. by mapping to reference sequence or PDB structures, and some contain VariO annotations.

The available datasets have been categorized into 20 groups and subgroups as indicated in Figure 2. The figure shows also the relationships of the datasets in different categories. Variants are often described at three molecular levels (DNA, RNA and protein) and sometimes also at protein structural level, including relevant cross references and variant descriptions. VariBench utilizes and follows a number of standards and systematics including HGVS variation nomenclature, HGNC gene names (not in all databases due to mapping problems), and VariO annotations in some datasets.

**Figure 2.**
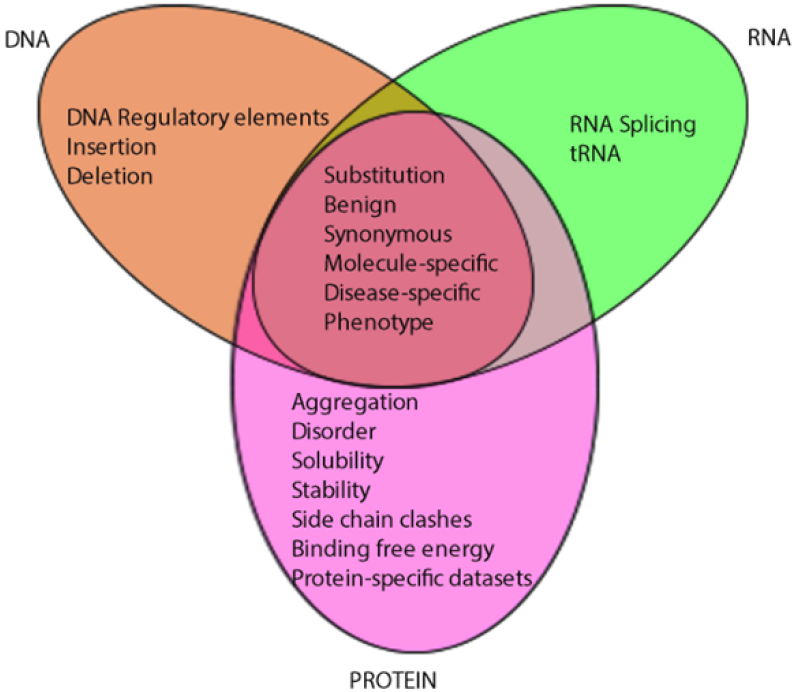
Types of benchmark datasets and their relations in VariBench.

Links are available to data in some external databases, including AmyLoad [84] and WALTZ-DB [85] for protein aggregation, DBASS3 and DBASS5 [71, 72] for splicing variants, SKEMPI [86], cancer datasets in KinMutBase [110], Kin-Driver [111], dbCPM [128], DoCM [130], and OncoKB [129], and tolerance predictor training set in DANN [47]. The latter has a link due to its huge size, the others since they are databases and as such easy to use directly and updated by third parties. We excluded datasets used in CAGI experiments, since they are available for registered participants only. LSDBs were excluded because data from these sources usually have to be manually selected before using as benchmark. Most of the time, there is no clear information for variant relevance to disease(s). Datasets for structural genomic variants were excluded, because they usually lack information about exact variation positions.

Unfortunately, many papers, even those reporting on benchmarking, do not contain and share the data, which does not allow others to extend the analyses and reuse the datasets.

### Variation type datasets

Variation types include insertions and deletions, coding and non-coding region substitutions, which are divided into training and test datasets, structure mapped variants, as well as synonymous, and benign variants. There are now data from four amino acid insertion effect predictors, mainly for short alterations. Only datasets added after the release of the first version of VariBench are discussed here. In Table 1 it is shown how many datasets and publications in each category appeared in the first edition.

Training datasets have mainly been used for development of machine learning predictors, there are 17 new datasets. They typically also contain test sets. Six test datasets have been specifically designed for method assessments. These include a set for addressing circularity [14] and pathogenicity/tolerance method performance assessment [57]. The American College of Medical Genetics and Genomics (ACMG) and the Association for Molecular Pathology (AMP) has published guidelines for variant interpretation [131]. These include instructions for use of prediction methods. A dataset was obtained for addressing concordance of prediction methods [56]. Another study addressed discordant cases [58]. Protein sequences of even closely related organisms contain differences and some of these are compensated variants where a disease-related variant in human is normal in another organism due to additional alteration(s) at other site(s). A dataset has been collected for such variants [59]. Unfortunately only the benign variants were made available. Analysis of the dataset representativeness, how well the datasets represent the variation space, was investigated for 24 datasets in VariBench and VariSNP [6]. These cases were mapped to reference sequence and are now available in the database.

Variations are mapped into protein three-dimensional structures in several datasets. Dedicated datasets contain those used for developing a method for predicting side chain clashes due to residue substitutions [60], analysis of effects on structures and functions of substitutions [61], and investigation of variations in membrane proteins [19].

There are two datasets for synonymous variants as well as two for benign ones.

### Effect-specific datasets

These datasets are for various types of effects. On DNA level there are 8 sets for DNA regulatory elements, and on RNA level 13 datasets for splicing. Most of the splicing datasets are very small, but there are a few with substantially larger numbers. In the first version of VariBench, there were only protein stability datasets in this category, totally 6 datasets from 4 studies. Thus the growth has been substantial.

Many more sets are available for effects on protein level. Protein aggregation (2 datasets), binding free energy (2), disorder (1), solubility (1), and stability are the currently available categories. Among protein stability datasets, there are 18 new datasets for single variants, almost all originating from ProTherm, and one dataset for double variants.

### Molecule-specific datasets

Thera are in VariBench 17 specific datasets for certain molecules. There is a set of variants used to train PON-mt-tRNA for substitutions affecting mitochondrial tRNA molecules [107]. This is of special interest as there are 22 unique mitochondrial tRNAs, which are implicated in a number of diseases.

The other datasets are protein specific. Kinact [109], Kin-Driver [111], KinMutBase [110], Kin-Mut [114] and the protein kinase dataset [112] contain variation information for protein kinases. The PON-BTK dataset was used to train a predictor for kinase domain variants in Bruton tyrosine kinase (BTK) [108]. There is a set for mismatch repair (MMR) proteins MLH1, MSH2, MSH6 and PMS2 and used to train PON-MMR2 [106].

Single amino acid substitutions were collected in 82 proteins to test whether there is a difference in performance for protein specific and generic predictors [12]. All the datasets contain at least ~100 variants. The results indicated vast differences in performances, the best generic predictors outperforming the specific predictors in most but not all cases.

The remaining datasets in this category are for variants in individual genes/proteins.

### Disease-specific datasets

This category contains totally 9 datasets, six of which are for cancer, one for long QT syndrome [125] and another for hypertrophic cardiomyopathy [126].

Although there are numerous studies of cancer variations, the functional verification of the relevance of those variants for the disease is usually missing. VariBench contains three datasets for variants in cancer, which have been experimentally tested [122–124], and links to three other sources, namely dbCPM [128], DoCM [130], and OncoKB [129]. In addition, there is the FASMIC dataset for variants which are largely cancer related [127].

### Phenotype dataset

One dataset contains information for disease phenotype, whether there is mild/moderate or severe disease due to substitutions. This dataset was used to train disease severity predictor called PON-PS [36].

## BENCHMARK USE CASE

VariBench datasets have mainly been used for prediction method development and testing. As the benchmark studies typically have not contained all the best performing tools, we compared the performance of the variant tolerance/pathogenicity predictor PON-P2, since this tool has been the best or among the best performing methods in a number of previous investigations [10, 12, 18, 19, 58]. The setup was similar in all these studies: to test the outcome of a spectrum of methods. We extended the published benchmark studies by repeating the original analyses with PON-P2. To avoid circularity, we first excluded from the datasets all cases that had been used for training PON-P2. The results are shown in Table 2 and are reported according to the published guidelines [32] and including some additional measures.

**TABLE 2.** Performance of PON-P2 on test datasets

| **Dataset** | **TP** | **FP** | **TN** | **FN** | **Coverage** | **PPV** | **NPV** | **Sens** | **Spec** | **Acc<sup>a</sup>** | **MCC** | **OPM** |
| --- | --- | --- | --- | --- | --- | --- | --- | --- | --- | --- | --- | --- |
| MutationTaster2, ClinVar [5] | 544 | 9 | 959 | 32 | 0.685 | 0.99 | 0.947 | 0.944 | 0.991 | 0.968 | 0.936 | 0.910 |
| MutationTaster2 [5] | 407 | 10 | 803 | 63 | 0.635 | 0.986 | 0.881 | 0.866 | 0.988 | 0.927 | 0.860 | 0.810 |
| Circularity, PredictSNPSelected [14] | 5116 | 341 | 3173 | 590 | 0.623 | 0.940 | 0.770 | 0.900 | 0.860 | 0.880 | 0.730 | 0.606 |
| Circularity, SwissVarSelected [14] | 1551 | 818 | 3194 | 773 | 0.557 | 0.650 | 0.810 | 0.670 | 0.800 | 0.750 | 0.460 | 0.325 |
| ACMG/AMP, MetaSVM [56] | 2588 | 364 | 2457 | 192 | 0.503 | 0.878 | 0.927 | 0.931 | 0.871 | 0.901 | 0.803 | 0.733 |
| ACMG/AMP, ClinVar_balanced [56] | 841 | 136 | 608 | 69 | 0.455 | 0.835 | 0.915 | 0.924 | 0.817 | 0.871 | 0.746 | 0.666 |
| ACMG/AMP, VaribenchSelected_Tolerance [56] | 1727 | 171 | 2996 | 57 | 0.513 | 0.947 | 0.967 | 0.968 | 0.946 | 0.957 | 0.914 | 0.875 |
| ACMG/AMP, predictSNPdsel [56] | 3752 | 317 | 3071 | 427 | 0.539 | 0.906 | 0.899 | 0.898 | 0.906 | 0.902 | 0.804 | 0.734 |
| ACMG/AMP, ClinVar_Sep2016 [56] | 1050 | 215 | 1726 | 102 | 0.514 | 0.892 | 0.909 | 0.911 | 0.889 | 0.900 | 0.801 | 0.729 |
| ACMG/AMP, Dominant_Recessive_Genes [56] | 1284 | 98 | 619 | 52 | 0.506 | 0.875 | 0.957 | 0.961 | 0.863 | 0.912 | 0.828 | 0.769 |
| ACMG/AMP, Oncogenes_TSG [56] | 535 | 59 | 74 | 3 | 0.497 | 0.692 | 0.99 | 0.994 | 0.556 | 0.908<br>0.775(AN) | 0.613 | 0.559 |
| Variants in 3D structures [46] | 5077 | 300 | 1060 | 266 | 0.337 | 0.812 | 0.94 | 0.95 | 0.779 | 0.865 | 0.741 | 0.676 |
| ClinVar dataset [57] | 1040 | 157 | 1200 | 169 | 0.541 | 0.881 | 0.864 | 0.86 | 0.884 | 0.872 | 0.745 | 0.664 |
| TP53 dataset [57] | 430 | 130 | 13 | 3 | 0.509 | 0.522 | 0.929 | 0.993 | 0.091 | 0.769<br>0.542(AN) | 0.195 | 0.269 |
| PPARG dataset [57] | 131 | 1376 | 7 | 0 | 0.598 | 0.501 | 1.000 | 1.000 | 0.005 | 0.503 | 0.000 | 0.111 |
| Cancer, functionally tested [123] | 561 | 18 | 16 | 3 | 0.605 | 0.653 | 0.989 | 0.995 | 0.471 | 0.965<br>0.733(AN) | 0.546 | 0.523 |
| Cancer, non-COSMIC functionally tested [123] | 108 | 10 | 14 | 3 | 0.455 | 0.700 | 0.956 | 0.973 | 0.583 | 0.904<br>0.778(AN) | 0.604 | 0.549 |

<sup>a</sup>AN, after normalization.

The exercise indicated that reproducibility and reusability could not be achieved in a number of cases due to problems in reporting. We had to exclude some published benchmark studies. The dataset for pharmacogenetics variants [54] was too small for reliable estimation. The paper for compensated variants [59] did not share the disease-related variants, and thus could not be evaluated. Of the dataset used by [53] only 36 cases were not included to the PON-P2 training set, and therefore had to be excluded.

We were able to perform the analysis for six studies and we analysed altogether 17 datasets. Full comparison was not possible in all cases as some details were not available. Therefore we discuss and compare the performances based on the information in the original papers, but list all the details from our study in Table 2.

For MutationTaster2 the published test data has not been previously available due to an inappropriate distribution format. MutationTaster 2 was originally compared to five tools and versions (MutationTaster1, PolyPhen humdiv and humvar, PROVEAN and SIFT) [5]. The accuracy and specificity are better for PON-P2 than the scores for the six tested tools and sensitivity is the second best. Only the measures given in the original article are discussed in here.

The study of circularity problems in variant testing was conducted on predictSNPSelected and SwissVarSelected datasets [14]. The performance of PON-P2 is superior compared to the eight tested predictors (MutationTaster2, PolyPhen, MutationAssessor, CADD, SIFT, LRT, FatHMM-U, FatHMM-W, Gerp++, and phyloP). In the test for predictSNPSelected dataset, NPV, PPV, sensitivity, accuracy and MCC are the best for PON-P2. Only for specificity it is the second best predictor, with a margin of 1%. In the data for SwissVarSelected, PON-P2 has the best score for PPV, accuracy and MCC. It is the second best for NPV and specificity, by 1-2% margin to the best, and for sensitivity. On both datasets, PON-P2 showed the most balanced performance.

25 tools were tested according to ACMG/AMP guidelines using several datasets [56]. The compared methods were REVEL, VEST3, MetaSVM, MetaLR, hEAt, Condel, MutPred, Mcap, Eigen, CADD, PolyPhen2, PROVEAN, SIFT, EA, MutationAssessor, MutationTaster, phyloP100way, FATHMM, DANN, LRT, SiPhy, phastConst100way, GenoCanyon, GERP, and Integrated_fitCons. Unfortunately, the results were not comprehensively reported. The paper contains data for AUC scores but they are presented as figures. The exact values were difficult to estimate, especially when results for 18 datasets were combined into single figures. In the end, we performed the test for 8 of these datasets. In the ClinVar balanced data the AUC of PON-P2 is either shared first or second, and in VariBenchselected data it has the best performance. Comparison for the six other datasets is not as reliable, but we can summarize that the PON-P2 performance is among the best if not best for all of these. It is really a pity that exact numbers were not provided by the authors.

The performances of 23 methods (FATHMM, fitCons, LRT, MutationAssessor, MutationTaster, PlyPhen humdiv and humvar versions, PROVEAN, SIFT, VEST3, GERP++, phastCons, phyloP, SiPhy, CADD, DANN, Eigen, FATHMM-MKL, GenoCanyon, M-CAP, MetaLR, MetaSVM, REVEL) were tested on three datasets: ClinVar and two protein specific sets for TP53 and PPARG [57]. They had also a fourth set for autism spectrum diseases, but since there is no experimental evidence for the relation of these variations to the disease that set was excluded. Although the study was well performed and described, it seems that the authors have not corrected for class imbalance. For the methods to be comparable the measures should be calculated based on the same data and have equal numbers of positive and negative cases. If that is not the case, the imbalance has to be mitigated with one of the available solutions. Some of the other benchmark studies may suffer from the same problem, but we are not sure due to incomplete descriptions of the studies. None of the tools can predict all possible variations and thus they have predictions for different numbers. Therefore we present the results both for non-normalized and normalized data. We believe that the former was used by the authors. In the case of ClinVar data, PON-P2 has better PPV, accuracy and MCC than the other methods tested in the paper.

In the case of TP53 data, the PON-P2 accuracy is second best when the data are not normalized, on other measures PON-P2 is ranked the fourth or worse. All cancer variants, such as those in TP53, were excluded from the PON-P2 training data. This was done because the effects of variations in cancers usually have not been experimentally verified. A variant in TP53 is not “pathogenic” alone, several variants in different proteins are needed for cancer.

All the predictors are known to have variable performance depending on the tested protein, see the study of protein-specific predictors [12]. That study showed that PON-P2 had better performance for 85% of proteins, being the best of the five tested tools (PolyPhen-2, SIFT, PON-P2, MutationTaster2, CADD). PPARG seems to be another example for which PON-P2 has poor performance [57]. An additional reason for poor performance may be that the PPARG data is not for pathogenicity, instead it is a “function score” that is based on the distribution of FACS sorted cells [132]. The same applies to the TP53 test data which is based on the protein function, not pathogenicity. Depending on a protein, the threshold for phenotype can be anything between 1 and 85% of the wild type activity (Vihinen, in preparation). We have previously tested PON-P2 in protein function prediction but with poor [133] or mixed (Kasak et al., submitted) outcome. This is because the method has not been trained and intended for this task. These results indicate the importance of applying computational tools to their intended purpose or at least testing the performance carefully before applied to new tasks.

Another study tested the performance of 14 tools (SEQ+DYN, SEQ, DYN, MutationTaster2, PolyPhen2, MutationAssessor, CADD, SIFT, LRT, FATHMM-U, Gerp++, phyloP, Condel, Logit) in relation to structural dynamics, which was used as a proxy for functional significance of amino acid substitutions [46]. PON-P2 has the best sensitivity, specificity, NPV and MMC, it is the second best for accuracy but only 13^th^ for PPV. The explanation for the latter observation is that many of the tested tools are severely biased, having very high PPV but very low NPV, whereas the performance of PON-P2 was again balanced over all the measures.

The exercise indicated that it is possible to compare predictors to published results based on exactly the same datasets. The new performance results for PON-P2 are in line with several previously published studies that have indicated the methods to be a top performer on different benchmarks [10, 12, 18, 19, 58]. When choosing a method(s), one should look at consistent performance over several benchmarks.

Full comparisons were not always possible because of incomplete performance assessments. Therefore, authors should meticulously describe all details and procedures in the data analysis as well as share the datasets used. Even if the data is taken from public sources, it is not possible for others to obtain exactly the same dataset as used in the papers even when applying the same selection criteria, as some important aspects seem always to be missing. In summary, it was possible to compare performances for methods not included into original studies. This is important in many ways, and contributes towards increased reproducibility and comparability. Good datasets are difficult to obtain, therefore VariBench will serve as a hub for sharing these important data.

